# Sleep is associated with reduction of excitatory signaling in medial prefrontal cortex

**DOI:** 10.1101/2025.04.29.651178

**Authors:** Takashi Yamada, Aaron Cochrane, Theodore LaBonte-Clark, Takeo Watanabe, Yuka Sasaki

## Abstract

Although many sleep medications enhance inhibitory signaling, it remains unclear whether inhibitory or excitatory neurotransmitters contribute to the natural transition from wakefulness to sleep in humans. Here, we show that changes in excitatory, rather than inhibitory, neurotransmitter levels are associated with this transition. Young, healthy participants underwent two nap sessions with polysomnography, during which glutamate and GABA concentrations in the medial prefrontal cortex were measured using magnetic resonance spectroscopy. Glutamate gradually decreased during deeper sleep stages compared to wakefulness in the second session, with better sleep quality. No such change occurred in the first session with poorer sleep, likely due to the first-night effect. Furthermore, reduced glutamate significantly mediated sleep-onset latency in both sessions. Conversely, GABA concentration did not change from wakefulness to sleep in either session. These findings provide the first evidence that reduced excitatory signaling is a key feature of natural good sleep onset in the human brain.

## INTRODUCTION

Understanding the neural mechanism of the transition from wakefulness to sleep is one of the most important issues in neuroscience. However, the mechanism underlying this transition is not yet fully understood in the human brain. Unlike many other animals, humans typically consolidate their sleep into a single, extended period each night once per night. This consolidation typically leads to high sleep quality and a short sleep onset latency (SOL), which is the time it takes to fall asleep after intending to do so. Prolonged SOL and low sleep quality are among the most critical issues in sleep problems including insomnia^1,2^. Therefore, addressing how SOL is regulated and how the gradual transition to deeper sleep occurs is essential not only for advancing our understanding of human sleep mechanisms but also for improving human sleep and cognition, as sleep is closely associated with human cognition^3,4^.

The neural mechanisms underlying sleep-wake regulations have been extensively investigated in animal models. It is well accepted that the reticular ascending arousal system^5^ or awake-promoting systems^6-10^ plays an important role in determining SOL and the transition to deeper sleep, as the suppression of these awake-promoting systems induces sleep in animals. The awake-promoting systems are assumed to span from subcortical regions, where numerous awake-promoting nuclei have been identified, projecting to cortical regions. Recent animal studies have highlighted the crucial roles of both glutamate and gamma-aminobutyric acid (GABA) within these awake-promoting systems^6-8,10^.

However, the precise neurochemical dynamics in the human brain during the wake-sleep transition remain largely elusive. Since many sleep medicines work through GABA receptors, the general public may assume that GABA release plays a critical role in promoting sleep onset and supporting the gradual transition to deeper sleep^11^. This assumption may lead to the idea that the inhibitory neurotransmitter concentrations increase during the transition. However, considering the involvement of both GABA and glutamate in awake-promoting systems, it is plausible that the concentrations of both inhibitory and excitatory neurotransmitters undergo changes in the human cortex during this transition.

The present study aimed to evaluate whether the concentrations of glutamate and GABA change in the human brain in a sleep-state-dependent manner. To capture these neurotransmitter dynamics, we repeatedly measured neurotransmitter concentrations during sleep using magnetic resonance spectroscopy (MRS) throughout the nap sessions while simultaneously polysomnography (PSG) was used to score sleep stages^12-14^. Repeated MRS measurements^15-18^ allowed us to track changes in the neurotransmitter concentrations across different states of wakefulness and sleep, includig the transition to deeper sleep in humans. In particular, we targeted the medial prefrontal cortex (mPFC) for neurotransmitter measurement, because the mPFC is recognized as a cortical region receiving projections from subcortical components of awake-promoting pathways in both animal models^6,7,10,19^ and humans^20,21^. In addition, the mPFC may be an important hub between the subcortical and cortical regions for regulating the level of consciousness and arousal^20^.

Moreover, we aimed to understand how changes in these neurotransmitter concentrations are associated with sleep quality. To achieve this, we employed a within-subject comparison of conditions characterized by varying SOLs, reflecting lower or higher sleep quality. For this purpose, we utilized the first-night effect (FNE)^22-31^. The FNE refers to a transient sleep disturbance occurring specifically during the first session of sleep experiments with healthy adults, often manifesting as prolonged SOL and lower sleep quality. This effect is known to occur not only during night sleep but also during daytime nap sessions^27-29^. Typically, the FNE diminishes by the second session of the sleep experiment^22-24,32^. By leveraging the FNE, we induced variations in sleep quality, as reflected by SOL, within the same participants, following established methodologies^25,30,33,34^. Therefore, we anticipated transiently disturbed sleep in the first session and improved sleep in the second session after a sufficient interval. To examine how neurotransmitter changes relate to sleep quality, we compared neurotransmitter concentrations between the first and second sleep sessions within the same subjects.

## RESULTS

### Confirmation of the FNE on Day 1

Following careful screening (see Methods for more details), twenty young and healthy participants without sleep complaints completed two early afternoon nap sessions (Day 1 and Day 2) in the early afternoon, separated by approximately one week according to our established sleep study protocol^12,13,28,35^. During both sessions, participants were instructed to lie down in the magnetic resonance imaging (MRI) scanner while concurrent polysomnography (PSG) was used for objective sleep stage scoring^36^.

We expected the sleep quality to be lower on Day 1 than Day 2 due to the FNE ^22-30^. To confirm the presence of FNE on Day 1, we examined whether the SOL, a metric known to be sensitive to this effect, and compared it between the two sessions. We predicted that if the FNE was present, SOL would be longer on Day 1.

As the distribution of our SOL data did not meet the assumption of normality, we used a nonparametric related-samples Wilcoxon signed-rank test to compare SOL between Day 1 and Day 2. The Wilcoxon signed-rank test revealed a significant difference in SOL between Day 1 and Day 2 (Z = -2.672, p = 0.008, n = 20), with Day 1 showing a significantly longer SOL (Figure 1). This result confirm that the sleep quality was lower on Day 1 compared to Day 2 due to the FNE on Day 1, consistent with recent literature^23,29^. Sleep architecture details are provided in Supplementary Table 1, and the number of participants reaching each sleep stage on Days 1 and 2 is shown in Supplementary Table 2.

**Table 1.** MRS data quality on Day 1 and 2.

|  | Day 1 |  |  | Day 2 |  |  | Wilcoxon signed-rank test |  |
| --- | --- | --- | --- | --- | --- | --- | --- | --- |
|  |  |  |  |  |  |  | Z | P |
| Shim values | 7.9 | ± | 0.65 | 7.0 | ± | 0.47 | -1.18 | 0.240 |
| %SD for GABA | 8.8 | ± | 0.50 | 9.4 | ± | 0.53 | 1.79 | 0.073 |
| %SD for Glx | 5.4 | ± | 0.16 | 5.5 | ± | 0.17 | 0.43 | 0.667 |
| NAA linewidth | 11.0 | ± | 0.48 | 10.1 | ± | 0.32 | -1.01 | 0.313 |
| Frequency drift | 2.3 | ± | 0.30 | 1.9 | ± | 0.22 | -1.27 | 0.204 |

Values are means ± SEM (n=20). The Wilcoxon signed-rank test was applied to test whether MRS data quality factors were significantly different between sleep sessions (Days 1 vs. 2). CRLB (%SD) represents the Cramer–Rao lower bounds, a measure of fitting error for metabolites. The NAA linewidth and frequency drifts are also commonly used to indicate the quality of MRS data.

**Figure 1.**
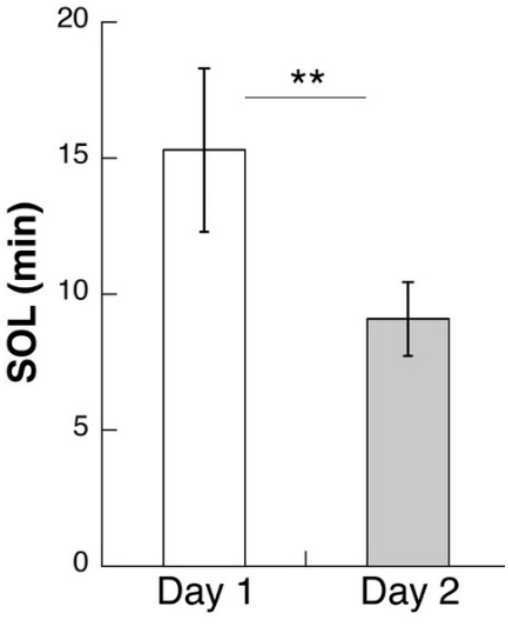
Sleep onset latency (SOL) in minutes. SOL was significantly longer on Day 1 compared to Day 2.

### Neurotransmitter changes during sleep

Following the co-registration of MRS and PSG data^12,13,29^, we obtained the mean concentrations of neurotransmitters, including Glx and GABA, for wakefulness, NREM sleep, and REM sleep. These data were obtained from the mPFC (Figure 2) for Day 1 and Day 2 (see Methods for further details). We confirmed a high degree of overlap (approximately 87%) in the voxel of interest (VOI) location between the two nap sessions (see Methods).

**Figure 2.**
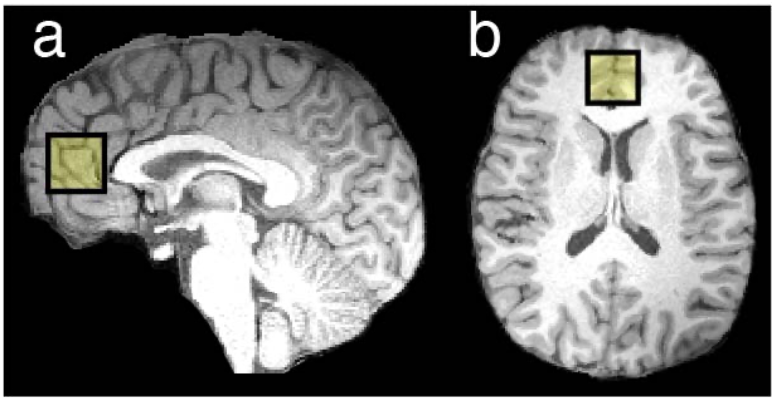
Voxel placement. VOI overlaid on structural images from a representative participant. a. Sagittal view. b. Transverse view.

We then calculated the percentage change in the neurotransmitter levels during sleep relative to a baseline defined as the average neurotransmitter levels during wakefulness. For example, the percentage change in Glx level during NREM sleep was calculated for each day using the following formula: [(Glx_NREM_ - Glx_wakefulness_)/Glx_wakefulness_ x 100 (%)] for each Day. In the present paper, we will refer to this percentage change (%) of neurotransmitter concentration as a ‘level’.

The subsequent sections will present the findings regarding Glx and GABA levels in the mPFC during NREM sleep. It is important to note that the data distributions for both Glx and GABA did not satisfy the normality assumption. Consequently, we primarily employed nonparametric statistical tests. Due to the limited number of participants who reached REM sleep in both sessions (Day 1 and Day 2, see Supplementary Table 2), we have not included the analysis of Glx and GABA levels during REM sleep in this study.

#### Glx levels during sleep stages

We tested whether the Glx level in the mPFC during NREM sleep significantly differed from the wakefulness baseline, using a one-sample Wilcoxon signed-rank test, performed separately for each Day. On Day 1, after correcting for two comparisons (Day 1 and Day 2), the one-sample Wilcoxon signed-rank test indicated that Glx levels during NREM sleep did not significantly differ from the wakefulness baseline (Figure 3; Z = -1.05, FDR-corrected p = 0.296, n = 20). In contrast, on Day 2, the one-sample Wilcoxon signed-rank test revealed a significant difference in Glx levels during NREM sleep compared to the baseline (Figure 3; Z = -3.51, FDR-corrected p < 0.001, n = 20). Furthermore, a related-samples Wilcoxon signed-rank test demonstrated a significant difference in Glx levels during NREM sleep between Day 1 and Day 2 (Z = -2.43, p = 0.015, n = 20, Figure 3).

**Figure 3.**
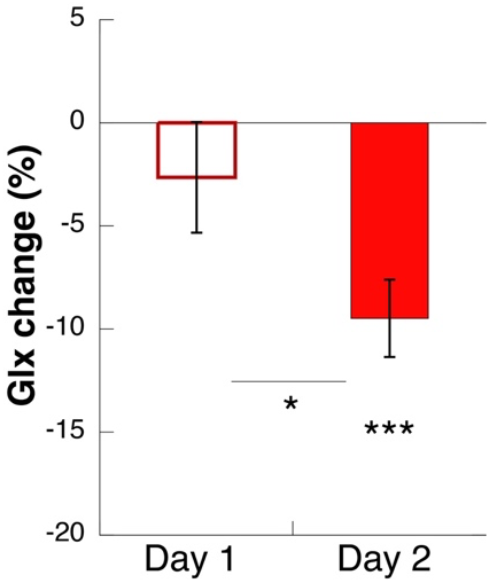
Glx levels during NREM sleep. The mean change (± SEM) in Glx levels during NREM sleep from the wakefulness baseline is shown for Day 1 and Day 2. Significance levels are indicated as * for p < 0.05 and *** for p < 0.001.

To examine the dynamics of Glx levels during the progression through sleep stages, we measured Glx levels in each NREM sleep stage (N1, N2, and N3). Given that N2 is frequently considered the stage of sleep onset in healthy individuals^12^, a significant change in Glx levels during N2 compared to wakefulness would indicate an association between Glx and sleep onset. Moreover, increasingly pronounced changes in Glx levels relative to wakefulness as sleep deepens (from N1 to N3) would suggest that Glx levels are sensitive to the gradual transition to deeper sleep. We used one-sample Wilcoxon signed-rank tests to determine whether Glx levels during each NREM sleep stage (N1, N2, and N3) significantly differed from the wakefulness baseline (represented as zero in our relative change calculation). The amount of MRS data available for each sleep stage is detailed in Supplementary Table 3, as not all participants exhibited all NREM sleep stages during the nap sessions.

On Day 2, when sleep quality was improved compared to Day 1 (Figure 4, filled red circles), we found significant decreases in Glx levels from the wakefulness baseline (zero) during N2 (Z = -2.20, FDR-corrected p = 0.042, n = 20) and N3 (Z = -2.67, FDR-corrected p = 0.024, n = 9), but not during N1 (Z = -1.05, FDR-corrected p = 0.296, n = 20), following FDR correction for three comparisons. These findings suggest that on Day 2, associated with improved sleep quality, a significant reduction in Glx levels occurred both at sleep onset (N2) and during the transition to deeper sleep (N3).

**Figure 4.**
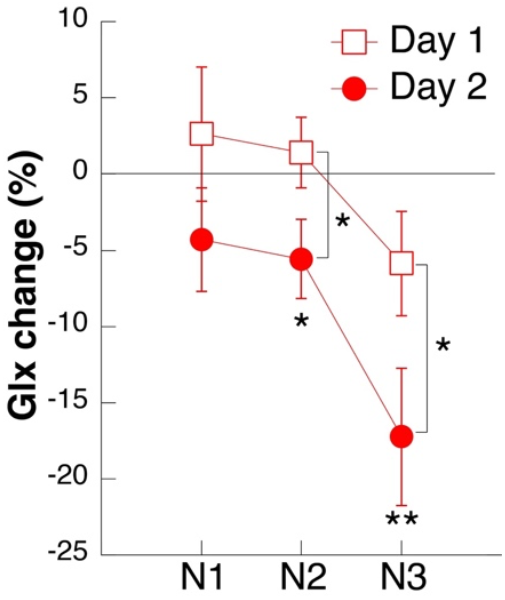
Glx levels during NREM sleep. The mean Glx level (± SEM) is shown for each NREM sleep stage (N1, N2, and N3). Squares represent Day 1, and circles represent Day 2. Significance levels are indicated as * for p < 0.05 and ** for p < 0.01.

A similar analysis was performed to examine Glx levels during N1, N2, and N3 on Day 1, which was characterized by lower sleep quality (Figure 4, squares). The one-sample Wilcoxon signed-rank tests showed no significant changes in Glx levels during any of the NREM sleep stages on Day 1 (N1: Z = 0.59, p = 0.554, n = 17; N2: Z = 0.11, p = 0.911, n = 20; N3: Z = -1.51, p = 0.131, n = 11) without correction for multiple comparisons.

Finally, we investigated whether the Glx level reductions during N2 and N3 on Day 2 were significantly different from those on Day 1. Related-samples Wilcoxon signed-rank tests, with FDR correction applied across the N2 and N3 stages, confirmed significant differences in Glx levels between Day 1 and Day 2 for both stages (N2: Z = -2.24, n = 20, FDR-corrected p = 0.046; N3: Z = -1.99, n = 6, FDR-corrected p = 0.046). These results indicate a significant reduction in Glx levels in the mPFC during the improved sleep condition on Day 2. Moreover, the results suggest that the Glx reduction was greater in the deeper sleep stages.

#### Glx changes during sleep and sleep quality

The preceding results suggest an association between a significant reduction in Glx levels and both improved sleep quality (as observed on Day 2 compared to Day 1) and deeper sleep stages. Therefore, we next investigated the relationship between Glx levels during NREM sleep and SOL, which we used as an indicator of sleep quality. We found a significant rank correlation between the Glx level in the mPFC during NREM sleep and SOL both on Day 1 and Day 2 (Supplementary Figures 1a, b; Spearman’s rho = 0.60, FDR corrected p = 0.010, n = 20 for Day 1; Spearman’s rho = 0.53, FDR corrected p = 0.017, n = 20 for Day 2) after correction for two comparisons.

These results suggest a close link between shorter SOL and a more pronounced Glx reduction in the mPFC during sleep. On Day 2, characterized by shorter SOL and improved sleep quality compared to Day 1 (Figure 1), we observed a significant Glx reduction during NREM sleep, whereas on Day 1, the FNE resulted in longer SOL, and no significant Glx reduction occurred during NREM sleep (Figure 3). This suggests a close association between the experimental day (reflecting sleep quality), Glx levels, and SOL. However, the longer SOL observed on Day 1 compared to Day 2 is a well-established effect of the FNE^23,27-29,31^. This raises the question of whether the observed Glx reduction is truly associated with SOL, or simply a byproduct of the different experimental sessions (Day 1 vs. Day 2).

To address this question, we employed a mediation analysis^37,38^ to investigate whether Glx levels during NREM sleep significantly mediated the effect of Day (Day 1 vs. Day 2) on SOL. In the mediation analysis (see Statistical analysis for details), we defined ‘Day’ as the independent variable, Glx level as the mediator, and SOL as the dependent variable. Prior to the analysis, both Glx levels and SOL were normalized using the Yeo-Johnson transformation^39^. The mediation analysis decomposes the total effect into a direct effect and an indirect (mediating) effect. First, we regressed SOL on Day to determine the total effect of Day on SOL, which was significant (Figure 5, path c, unstandardized coefficient = -4.19, p = 0.001, n = 20), indicating a significant association between Day and SOL. Second, we investigated the relationship among these three factors, using structural equation modeling^38^ for mediation analysis. The results indicated a significant effect of Day on Glx levels (Figure 5, path a, unstandardized coefficient = -6.14, p = 0.004, n = 20), and subsequently, a significant effect of Glx levels on SOL (Figure 5, path b, unstandardized coefficient = 0.34, p < 0.001, n = 20). Further, after accounting for the mediating effect of Glx levels, the direct effect of Day on SOL was no longer significant (Figure 5, path c’, unstandardized coefficient = -2.09, p = 0.106, n = 20), suggesting a full mediation of the effect of Day on SOL by Glx levels. Finally, the mediation effect of Glx levels on SOL (Figure 5, a x b, unstandardized coefficient = -2.20, p = 0.028, n = 20) was significant, accounting for approximately 51% of the total effect of Day on SOL. These results indicate that the Glx level during NREM sleep significantly mediated the effect of Day on SOL.

**Figure 5.**
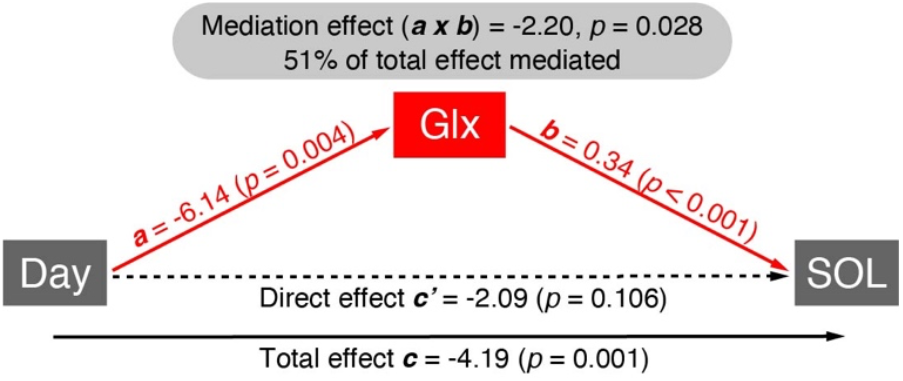
Glx levels during NREM sleep mediate SOL. Our tripartite model investigated the relationship between Day, SOL, and Glx levels during NREM sleep. Specifically, we explored how Day impacts SOL through changes in Glx. The total effect (path c, black solid line) is the sum of direct effect (path c’, black dashed line) and indirect or mediating effect (red lines, a x b). The proportion of the total effect mediated by Glx (a x b / (a x b + c’)) was 0.513 [(−2.2) / (−2.2 + (−2.09))]. See the main text for statistical details.

#### GABA levels during sleep stages

Next, we assessed whether the GABA level in the mPFC during NREM sleep significantly differed from the wakefulness baseline. Although the overall trend showed GABA levels slightly higher than baseline on Day 1 and slightly lower on Day 2, a one-sample Wilcoxon signed-rank test indicated no significant difference in GABA levels during NREM sleep from baseline on either Day 1 (Figure 6; Z = 1.61, p = 0.11, n = 20) or Day 2 (Z = -1.31, p = 0.191, n = 20) without correction for multiple comparisons. Moreover, a related-samples Wilcoxon signed-rank test revealed no significant difference in GABA levels between Day 1 and Day 2 (Z = -1.49, p = 0.135, n = 20). These results indicate no statistically significant change in GABA levels during NREM sleep compared to wakefulness, regardless of sleep quality.

**Figure 6.**
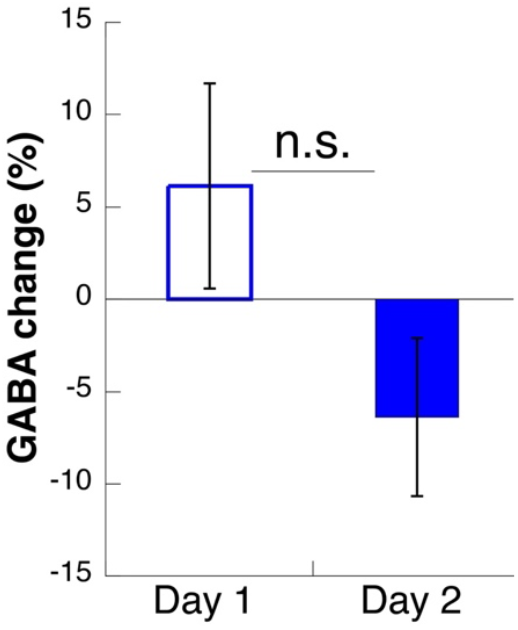
GABA levels during NREM sleep. The mean change (± SEM) in GABA levels during NREM sleep from the wakefulness baseline is shown for Day 1 and Day 2.

However, examining GABA levels across individual NREM sleep stages (N1, N2, and N3) suggested a potential elevation above baseline during lower-quality sleep. On Day 1, associated with lower sleep quality (Figure 7, squares), one-sample Wilcoxon signed-rank tests revealed significantly different GABA levels from baseline during N1 (Z = 2.911, FDR-corrected p = 0.012, n = 17) and N2 (Z = 2.203, FDR-corrected p = 0.0345, n = 20), but not during N3 (Z = 1.156, FDR-corrected p = 0.248, n = 11), following FDR correction for three comparisons. In contrast, on Day 2, when sleep quality was improved (Figure 7, filled circles), GABA levels did not significantly differ from baseline during N1 (Z = 0.635, p = 0.526, n = 20), N2 (Z = -0.336, p = 0.737, n = 20), or N3 (Z = -1.481, p = 0.139, n = 9) without correction for multiple comparisons. When comparing GABA levels for each sleep stage between Day 1 and Day 2 using related-samples Wilcoxon signed-rank tests, we found significant differences for N2 (Z = -2.128, FDR-corrected p = 0.0495, n = 20) and N3 (Z = -2.547, FDR-corrected p = 0.033, n = 6), but not for N1 (Z = -1.728, p = 0.084, n = 17), after FDR correction for three comparisons.

**Figure 7.**
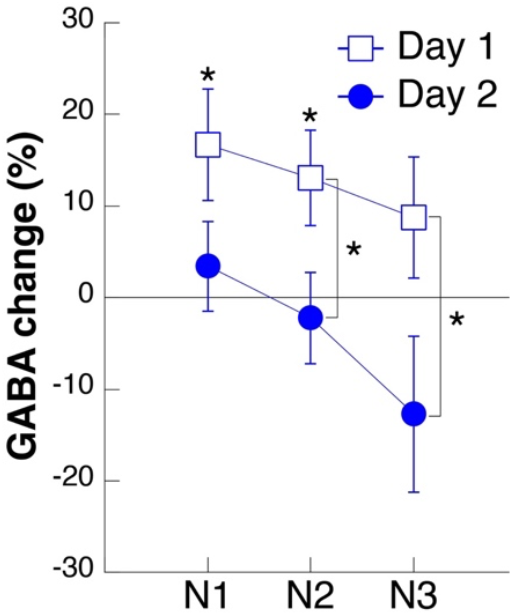
GABA levels during NREM sleep. The mean GABA level (± SEM) is shown for each NREM sleep stage (N1, N2, and N3). Squares represent Day 1, and blue circles represent Day 2. Significance levels are indicated as * for p < 0.05.

These results suggest that GABA levels remained stable relative to wakefulness during improved sleep on Day 2, whereas they appeared to increase during N1 and N2 under lower-quality sleep conditions on Day 1. Therefore, the association between GABA levels and sleep quality remains uncertain based on these findings.

#### GABA changes during sleep and sleep quality

We investigated the correlation between GABA levels during NREM sleep and SOL using a nonparametric Spearman correlation. In contrast to Glx levels, the GABA level did not significantly correlate with the SOL during NREM sleep on Day 1 (Supplementary Figure 2a, Spearman’s rho = -0.04, p = 0.870, n = 20) or Day 2 (Supplementary Figure 2b, Spearman’s rho = 0.12, p = 0.607, n = 20). These results indicate that GABA levels are not strongly associated with SOL in our data.

We conducted a mediation analysis to examine whether GABA levels during NREM sleep mediated the effect of Day (Day 1 vs. Day 2) on SOL. GABA levels and SOL were normalized using the Yeo-Johnson transformation^39^. Consistent with the Glx mediation analysis, we defined ‘Day’ as the independent variable, GABA level as the mediator, and SOL as the dependent variable. First, the total effect of Day on SOL was significant (Figure 8, black thick line, path c, unstandardized coefficient = -4.19, p = 0.001, n = 20). Next, structural equation modeling^38^ indicated a marginally significant effect of Day on GABA level (Figure 8, path a, unstandardized coefficient = -12.49, p = 0.065, n = 20), whereas the effect of GABA on SOL was not significant (Figure 8, path b, unstandardized coefficient = 0.06, p = 0.84, n = 20). Then, the mediation effect of GABA on SOL was not significant, either (Figure 8, mediation effect a x b, unstandardized coefficient = -0.073, p = 0.869, n = 20), with GABA levels accounting for less than 2% of the total effect of Day on SOL (Figure 8). Accordingly, even after accounting for the effect of GABA levels, the direct effect of Day on SOL remained significant (Figure 8, path c’, unstandardized coefficient = -4.12, p = 0.001, n = 20), confirming that GABA levels during NREM sleep did not significantly mediate the relationship between Day and SOL. These results imply that the role of GABA in sleep quality, as measured by SOL, is minimal.

**Figure 8.**
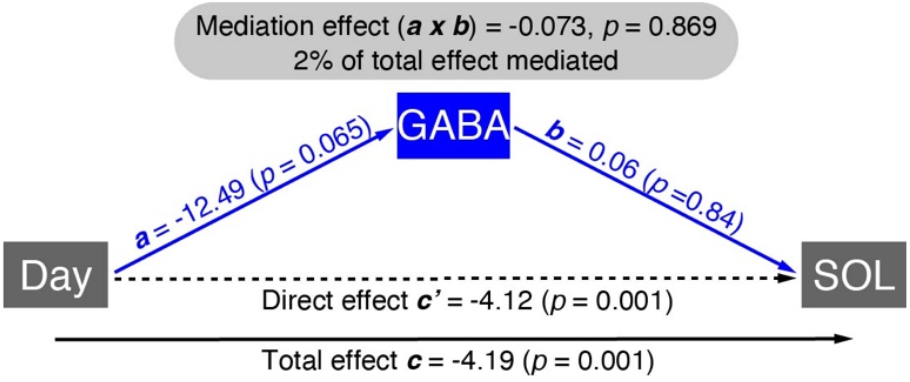
GABA levels during NREM sleep do not mediate SOL. Our tripartite model investigated the relationship between sleep sessions (Day), sleep onset latency (SOL), and GABA levels during NREM sleep. The total effect (path c, black solid line) is the sum of the direct effect (path c’, black dashed line) and the indirect or mediating effect (blue lines, a x b). The proportion of the total effect mediated by GABA (a x b / (a x b + c’)) was 0.017 [(−0.073) / (−0.073 + (−4.12))]. See the main text for statistical details.

### Control analyses

To ensure the robustness of our findings regarding Glx and GABA levels during NREM sleep on Day 1 and Day 2, we conducted three control analyses to rule out potential confounding effects of baseline neurotransmitter concentrations and MRS data quality.

First, we assessed whether N-acetylaspartate (NAA) concentrations, the control metabolite for Glx and GABA quantification, varied across sleep stages (wakefulness and NREM sleep) and Days. A related-samples Wilcoxon signed-rank test showed no significant difference in NAA levels between wakefulness (Figure 9a, white bars) and NREM sleep (Figure 9a, gray bars) on Day 1 (Z = 0.00, p = 1.000, n = 20) or Day 2 (Z = -1.68, p = 0.093, n = 20). Furthermore, NAA concentrations during wakefulness showed no significant difference between Day 1 and Day 2 (related-samples Wilcoxon signed-rank test; Z = 0.65, p = 0.514, n = 20). Similarly, NAA concentrations during NREM sleep showed no significant difference between Day 1 and Day 2 (Figure 9a, gray bars; Z = -0.30, p = 0.765, n = 20). These results indicate that the concentration of the control metabolite remained consistent across both sleep sessions (Day 1 and Day 2) and sleep stages (wakefulness and NREM sleep).

**Figure 9.**
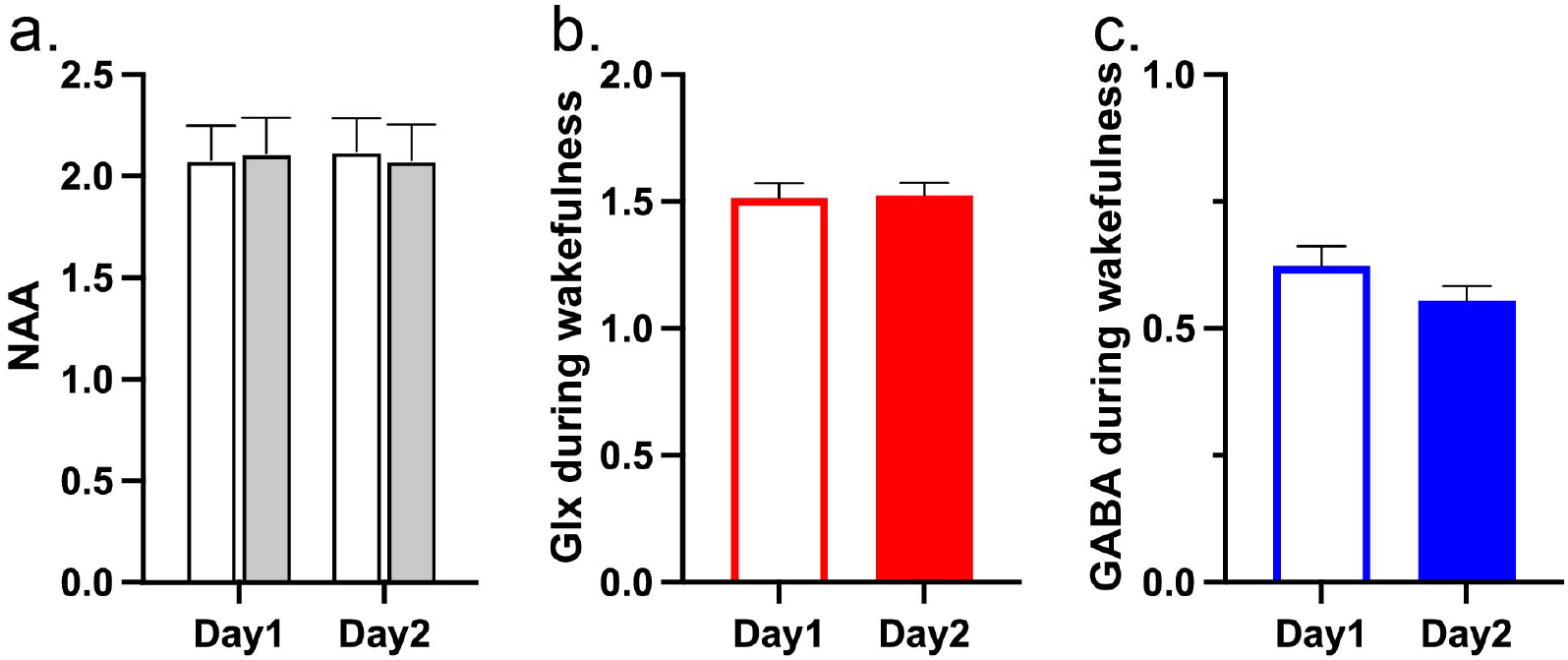
Concentrations of the control metabolite NAA and baseline concentrations of Glx and GABA. a. The mean absolute NAA concentrations (IU) (± SEM) during wakefulness (white bars) and NREM sleep (gray bars) on Day 1 and Day 2. b. The mean baseline Glx levels (Glx / NAA ratio) (± SEM) on Day 1 and Day 2. c. The mean baseline GABA levels (GABA / NAA ratio) (± SEM) on Day 1 and Day 2.

Second, we assessed whether baseline Glx and GABA concentrations during wakefulness differed between Day 1 and Day 2. Given that our primary analyses focused on the relative changes in Glx and GABA levels during NREM sleep compared to the wakefulness baseline within each session (Day 1 and Day 2), any significant difference in these baseline levels between sessions could potentially confound our interpretation. However, related-samples Wilcoxon signed-rank tests revealed no significant differences in wakefulness baseline concentrations of either Glx (Figure 9b; Z = 0.64, p = 0.526, n = 20) or GABA (Figure 9c; Z = -0.97, p = 0.332, n = 20) between Day 1 and Day 2. These findings indicate that baseline Glx and GABA levels during the wakefulness period were consistent across the two sleep sessions.

Third, we assessed whether the quality of MRS data differed significantly between the two sleep sessions. The quality of MRS data was assessed using four parameters: shim values, % SD, NAA linewidth, and frequency drift. We compared each of these parameters between the two sleep sessions. Statistical analyses revealed no significant differences in these parameters between the two sleep sessions (Table 1). These results indicate that the quality of MRS data was consistent across the two sleep sessions.

## DISCUSSION

A comprehensive understanding of the neural mechanisms underlying the wake-sleep transition in humans is still lacking. By simultaneously employing functional MRS and PSG, and leveraging the first-night effect (FNE), we successfully assessed neurotransmitter dynamics associated with sleep quality and the progression to deeper sleep stages in the human cortex. Notably, we observed that enhanced sleep quality and depth were marked by a significant reduction in excitatory neurotransmission, contrasting with the expectation of increased inhibitory activity.

We propose several potential explanations for the reduction in Glx levels observed across the wake-sleep transition and into deeper sleep. We speculate that this Glx reduction is closely linked to the deactivation of wake-promoting systems in the human brain, given the critical role of excitatory signaling in maintaining wakefulness^8,40,41^. Our Glx measurements were taken from the mPFC, a cortical region considered to be an integral component of wake-promoting circuits^8^. Furthermore, the mPFC serves as a crucial bidirectional hub connecting subcortical and cortical regions within awake-promoting systems^20,42-44^. Therefore, the observed reduction in excitatory signaling in the mPFC likely reflects the functional shift occurring within this key regulatory region during the transition to sleep.

A particularly important and somewhat surprising finding was the lack of a significant association between GABA concentration and sleep quality in our data. Several lines of evidence suggest a significant link between increased GABA concentration and improved sleep. First, it is well established that GABAergic neurons in subcortical regions promote sleep by inhibiting wake-promoting systems ^6-8,10,45^. Second, commonly prescribed hypnotic —including benzodiazepines and Z-drugs—are GABAergic. Contrary to this expectation, we observed no significant changes in GABA levels during sleep transitions on Day 2, which was characterized by improved sleep quality. In fact, a trend towards decreased, rather than increased, GABA levels was observed during the progression to deeper sleep stages on Day 2. Furthermore, GABA concentration did not mediate SOL. Collectively, these findings render the absence of a significant association between GABA levels and sleep quality particularly noteworthy and unexpected.

Several potential explanations could account for the absence of significant GABA modulation despite the improved sleep observed on Day 2. First, it is possible that increased GABA concentrations are indeed crucial for sleep promotion, occurring within specific inhibitory sleep-promoting circuits^45^, of which the mPFC may not be a key component. Consequently, other brain regions beyond the mPFC might exhibit increased GABA concentrations during the sleep transition. Second, the subtle increase in inhibitory signaling necessary for the suppression of wake-promoting systems might have been below the sensitivity limits of the MRS methodology employed in this study. Future studies utilizing higher magnetic field strengths and smaller voxel volumes could potentially detect such subtle GABAergic changes. However, the fact that we observed a tendency for GABA levels to be elevated during the poorer sleep quality on Day 1 (in light sleep) argues against a simple issue of GABA detectability with our MRS approach. Third, a more radical possibility is that GABA’s involvement in the human sleep transition is less direct than traditionally assumed. Despite the lack of a clear sleep-promoting role, the higher GABA concentrations observed on Day 1—during light sleep compared to wakefulness, and during N2/N3 compared to Day 2—suggest that GABA is not entirely inert during sleep transitions. One interpretation is that increased GABA levels might be associated with a less efficient or more effortful transition into deeper sleep. These findings suggest that GABA is not entirely irrelevant to sleep transitions. Further studies are needed to clarify the role of GABA in transitions from wakefulness to sleep and across sleep stages.

The current study focused on daytime nap sessions, and further research is necessary to ascertain the generalizability of these findings to nocturnal sleep. However, it is crucial to emphasize that we meticulously controlled for potential confounding effects of circadian timing in this study. Both the first and second nap sessions were scheduled at the same time of day, thereby minimizing the influence of circadian phase on the observed differences in sleep quality. Furthermore, the inter-session interval of over one week was designed to minimize any potential carryover effects or circadian rhythm disruption that might have been induced by the first nap session. Regular sleep-wake patterns in the participants prior to the nap sessions were verified through both objective actigraphy data and subjective sleep logs. Crucially, sleep-onset REM periods, a hallmark of circadian misalignment, were absent in all nap sessions ^31,46^ further supporting the controlled circadian context of our study. Therefore, it is highly improbable that the significant reduction in Glx concentration observed in the mPFC during the wake-sleep transition was a consequence of circadian timing disturbances.

We believe that our experimental design effectively isolated sleep quality as the primary differentiating factor between Day 1 and Day 2. First, by employing the FNE to manipulate sleep quality within a within-subject design, we minimized the influence of uncontrolled individual differences such as medication use and comorbidities. Second, we recruited healthy young adults without sleep complaints through a rigorous screening process. This approach reduced inter-subject heterogeneity, a common confound in studies comparing individuals with and without sleep disorders^47^. Third, consistent and precise voxel placement for MRS measurements across the two sessions minimized the likelihood that variations in MRS signals between Day 1 and Day 2 were attributable to differences in voxel locations. Fourth, the control analyses revealed no significant differences between Day 1 and Day 2 in baseline levels of Glx, GABA, the control metabolite NAA, nor in MRS data quality. Collectively, these findings strongly support the conclusion that the differential changes observed in Glx and GABA levels between the two nap sessions are highly attributable to the variations in sleep quality between Day 1 and Day 2.

In the present study, we focused on Glx and GABA. However, the significant roles of other neurotransmitters beyond glutamate and GABA in sleep-wake regulation are well documented^6-8,10,45,48^. Further studies are needed to elucidate the complex interactions between subcortical and cortical regions in humans, particularly concerning the dynamic interplay of various neurotransmitter systems during the transition from wakefulness to deeper sleep stages.

## METHODS

### Participants

Twenty-two young and healthy adults (13 females) aged 18–30 years participated in this study following careful screening (see below). Two participants were subsequently excluded from the analysis: one due to a failure to achieve sleep onset, and the other due to the absence of sustained wakefulness during the nap session, resulting in no usable data on neurotransmitter concentration changes from the wakefulness baseline to NREM sleep. Thus, the final analysis included data from 20 participants (mean age ± SD: 20.9 ± 2.9 years; 12 females). All participants provided written informed consent after receiving a thorough explanation of the study’s objectives and procedures. The study protocol was approved by the Institutional Review Board at Brown University.

Following informed consent, all participants completed the Morningness-Eveningness Questionnaire (MEQ)^49^ and the Pittsburgh Sleep Quality Index Questionnaire (PSQI)^50^. These assessments confirmed that all participants were intermediate chronotypes, with MEQ scores ranging from 42 to 69. Additionally, no participants were classified as poor sleepers, as indicated by PSQI scores of five or less.

### Screening

We conducted a careful screening process, consistent with our established methods to determine eligibility. This process excluded individuals with physical or psychiatric illnesses, those currently taking medication, or those with suspected sleep disorders based on self-reports^12,13,28,35^. Potential participants completed questionnaires, including a detailed sleep-wake habits questionnaire^12,13,28,29,35^ and the Munich Chronotype Questionnaire (MCTQ)^51^. These assessments enabled us to verify that participants maintained regular sleep-wake patterns, defined as average bedtimes and wake-up times with a consistent variation of no more than two hours between workdays and free days, and reported an average sleep duration of six to nine hours.

### Experimental design

To ensure consistent sleep quality across participants throughout the experimental sessions, which took place over a few weeks, we instructed them to maintain their regular sleep-wake habits, including stable sleep and wake times and durations, in the period leading up to the experiments. We also monitored their sleep-wake behavior for three to seven days prior to the sleep sessions using a sleep log and an actigraphy device (GT9X-BT, ActiGraph). If a participant’s sleep-wake schedule exhibited a variation of more than two hours between workdays and free days during this monitoring period, their sleep experiment was postponed until their sleep-wake schedule stabilized. Participants were also instructed to abstain from alcohol consumption, excessive physical activity, and napping starting from the day before the sleep sessions, due to their potential to negatively impact sleep quality. Participants were further instructed to avoid caffeine consumption on the day of the sleep sessions.

Participants underwent two early afternoon sessions, approximately one week apart, following our established sleep study protocols^12,13,28,35^. The at least one-week interval between sessions was implemented to ensure that any potential effects of the early afternoon nap in the first session did not carry over and influence sleep quality in the second session. The initial session (Day 1) was anticipated to exhibit lower sleep quality due to the FNE, compared to the second session (Day 2), which was expected to result in higher sleep quality.

In each session, prior to the nap, electrodes for PSG (see below) were attached to the participants outside the MRI scanner. Participants were then instructed to lie down inside the MRI scanner, during which time PSG data recording commenced, followed by the start of MRI acquisition (see below). The time window for the nap approximately corresponded to 2:30-4 pm. Additionally, the Stanford Sleepiness Scale (SSS)^52^ was administered to five participants immediately before each nap session to assess potential differences in subjective sleepiness between Day 1 and Day 2.

### Polysomnography (PSG)

PSG included electroencephalography (EEG), electrooculogram (EOG), electromyogram (EMG), and electrocardiogram (ECG). We used MRI-compatible EEG caps (Brain Products GmbH) for the PSG recordings (32-channel cap for 15 participants and 25-channel cap for 5 participants). The 32-electrode cap consisted of 23 EEG electrodes for the scalp, six for EOG, two for EMG, and one for ECG electrode. Horizontal EOGs were obtained from two bipolar electrodes positioned at the outer canthi of both eyes and vertical EOGs from four electrodes positioned above and below the left and right eyes. The 25-electrode configuration included 18 scalp EEG electrodes, four EOG electrodes, two EMG electrodes, and one ECG electrode. For this setup, EOGs were captured with bipolar electrodes at the outer canthi of each eye and two additional electrodes above and below the left eye.

Across both cap configurations, EMG was recorded bipolarly from the chin, while ECG was obtained from a site approximately two finger-widths left of the spine, at the level of the inferior scapulae. EEG and ECG electrodes were referenced to Fz, with the ground electrode positioned at AFp4. Electrode impedances were kept at or below ten kΩ for EEG and ECG and below 15 kΩ for EOG and EMG. Data acquisitions were performed using an MRI-compatible amplifier (BrainAmp MR, Brain Products) and recording software (BrainVision Recorder, Brain Products) at a sampling rate of 5,000 Hz.

Scanner and ballistocardiogram artifacts were removed using Brain Vision Analyzer 2 (Brain Products). Subsequently, the EEG data were re-referenced to the left (TP9) and right (TP10) mastoids.

### Sleep-stage scoring and sleep factors

Sleep stages were scored in every 30-second epoch according to the standard criteria defined in the American Academy of Sleep Medicine (AASM) Scoring Manual ^36^. The stages were as follows: wakefulness (stage W), NREM stages (N1, N2, and N3), and REM sleep (stage REM).

Sleep onset was defined by the first appearance of stage N2, consistent with previous research^12,13,27-29,35^. The time interval from lights-off to this sleep onset was defined as SOL.

See Supplementary Table 1 for the sleep structures and Supplementary Table 2 for the number of data available for each sleep stage.

### MRS acquisition

MRI data were acquired on a 3T Siemens Prisma scanner using a 64-channel head coil. To ensure participant comfort and minimize head movement during MRI measurements, their heads were securely positioned using cushions and gauze to prevent motion and discomfort. Additional support was provided upon request with back and knee cushions, and blankets were used to maintain body warmth and facilitate the onset of sleep.

For each sleep experiment, a T1-weighted anatomical reference image dataset was acquired using a magnetization-prepared rapid gradient echo (MP-RAGE) sequence to obtain high-resolution structural brain images. The imaging parameters were: repetition time (TR) = 1.9 s, echo time (TE) = 3.02 ms, flip angle (FA) = 9°, voxel size = 1 × 1 × 1 mm, 256 slices, and no slice gap. High-resolution brain images enabled precise localization of the voxel of interest (VOI) for subsequent MRS scans.

The voxel of interest (VOI) (Figure 2) was a 2.5 × 2.5 × 2.5 cm^3^ volume encompassing the mPFC, a size chosen to optimize the balance between spatial resolution and sufficient signal-to-noise ratio^53^. We manually placed the VOI anterior to the genu of the corpus callosum bilaterally based on the observed anatomical structure. The VOI was positioned to include the least white matter possible because its lipid content could potentially introduce noise into the MR spectrum. After the VOI placement, shimming, performed with an automated tool from the scanner’s manufacturer, optimized the magnetic field homogeneity around the VOI. The entire preparation process, including structural MRIs, VOI placement, and shimming, was completed within 20–30 minutes. We paid extra attention to place the VOI at the exact anatomical locations as possible across the first and second sleep sessions by referencing the structural images in high resolution. The position of the VOI demonstrated a high degree of consistency between Day 1 and Day 2, with an overall overlap of 86.6 ± 1.8% (mean ± SEM) in volume. The calculation of the overlap was done as follows. We aligned the structural data on Day 2 to those on Day 1 to obtain the degree of transformation, applied the same transformation to the Day 2 VOI, counted overlapping voxels between VOIs, and computed the percentage relative to the total VOI volume (15625 voxels) for each participant.

We employed the MEGA-PRESS sequence to quantify Glx (the combined concentration of glutamate and glutamine) and GABA within the VOI. This technique utilizes a frequency-selective editing pulse to isolate specific metabolite signals, consistent with established methodologies in previous research. The ‘Edit On’ and ‘Edit Off’ phases facilitated the differentiation of GABA and Glx concentrations from the composite spectrum. We first ran a shorter version of the MEGA-PRESS sequence (TR = 1.25 s; TE = 68 ms; FA = 90°; with 32 averages, lasting 2:05 minutes) to quickly test whether the MRS data quality was acceptable for the VOI placement. Cramer-Rao lower bounds (CRLB) were used to assess MRS data quality, and data were considered acceptable if the CRLB values for both Glx and GABA were below 25%, consistent with previous concurrent MRS-PSG studies (see Quality tests for the MRS data below)^12,13,29^. After confirming that the data quality was acceptable, we turned off the lights and started a longer MRS scan (TR = 1.25 s; TE = 68 ms; FA = 90°; with 240 averages, lasting 10:05 minutes), repeated 9 times.

It is important to note that we measured Glx, a composite of glutamate and glutamine, rather than isolated glutamate. This choice was made because it is uncertain whether linear-combination modeling can distinguish glutamate from glutamine and GABA in MEGA-PRESS sequence data^16^. Moreover, Glx (glutamate + glutamine) is considered a reliable measure^16^. However, the contribution of glutamate is estimated to be approximately 85% of Glx^54^. Thus, while we reported our results in terms of Glx, it is likely that our results predominantly reflect glutamate levels.

### Temporal co-registration of MRS and PSG data

Temporal co-registration of PSG and MRS data from the sleep sessions was performed according to procedures detailed in previous studies^12,13,29^. First, sleep was categorized into three stages: wakefulness, NREM sleep, and REM sleep. The NREM sleep category combined stages N1, N2, and N3. Initially, we divided the PSG and MRS data into 10-min blocks. Given that sleep stages were scored at 30-second intervals, each block included 20 epochs of sleep stages. To co-register MRS data to sleep stages, we applied a three-step process. First, if a single sleep stage was dominant and accounted for over 80% (i.e., 16 epochs) of a 10-minute block, we labeled the entire MRS block with that sleep stage. Second, if a single sleep stage did not dominate one block, we divided the 10-minute block into five 2-minute sub-blocks, each corresponding to four 30-second epochs. A sub-block was assigned to a sleep stage if it comprised more than 50% of that stage (i.e., two epochs). Third, in cases where a sub-block had equal amounts of two different stages, we prioritized the stages as follows: wakefulness, NREM sleep, and REM sleep. For example, a sub-block with an equal number of wakefulness and NREM stages would be labeled as wakefulness. If a sub-block contained all three stages, we assigned it to the stage with the most epochs. This system ensured that each block of MRS data reflected the predominant sleep stage. When we analyzed MRS levels for stages N1, N2, and N3 separately, we applied the same rules but expanded to five categories, prioritizing wakefulness followed by N1, N2, N3, and REM sleep. See Supplementary Table 3 for the number of data available for MRS data in each sleep stage.

### MRS analysis for Glx and GABA concentration during NREM sleep

We primarily quantified Glx, GABA, and NAA concentrations within the VOI using LCModel ^55,56^, focusing on the MRS signal within a specific chemical shift range (1.95 to 4.2 ppm). The LCModel operates under the assumption that the obtained spectrum can be fitted to a linear combination of spectra of individual metabolites, derived from an imported metabolite basis set (Supplementary Figure 3). We used NAA as a control metabolite to normalize the levels of Glx and GABA, following the procedure used in earlier research^12,15,29,57^. Corrections for water scaling and eddy current were carried out.

After the co-registration of MRS data with the PSG data in time, we quantified how much the concentrations of Glx and GABA changed in the transition from wakefulness to sleep. First, we calculated the average concentrations of each Glx and GABA during wakefulness and NREM sleep, respectively. The average concentrations of each neurotransmitter during wakefulness, which included the period from lights off (the start of the MRS measurement) to sleep onset and any instances of wakefulness after sleep onset, worked as the baseline. Second, we assessed the changes in these neurotransmitter levels during NREM sleep relative to baseline. For example, to calculate the relative change of Glx level during NREM sleep, we used this formula: [(Glx_NREM_ - Glx_wakefulness_)/Glx_wakefulness_ x 100 (%)].

### Quality tests for the MRS data

First, following previous studies, we used the CRLB to exclude low-quality MRS data ^12,29,58^. The CRLB indicate the reliability of the quantification of Glx, GABA, and NAA as measures of fitting errors. A threshold of 25% was used to reject low-quality MRS results^58-60^. This threshold rejected the average of 1.3% of all MRS data from two sleep sessions, with 1.5% rejected on Day 1 and 1.1% on Day 2.

Next, to compare MRS data quality between the two sleep sessions, we calculated four parameters for the spectra, as in previous studies: CRLB, shim values, NAA linewidth, and frequency drift^12,29^. The shim values represent the homogeneity of the magnetic field. NAA linewidth provides a measure of signal resolution, and frequency drift signifies the instability of the B0 magnetic field. We used these four indices to determine if there were differences in the signal quality of MRS data between Day 1 and Day 2 (Table 1).

### Statistical analysis

A two-tailed type I error rate (α-level) of 0.05 was used for all statistical tests. Data normality was assessed using the Shapiro-Wilk test. As most key measures violated the assumption of normality, we primarily employed nonparametric tests, including the Wilcoxon signed-rank test and Spearman’s correlation coefficient, with the exception of the mediation analysis (see below). The False Discovery Rate (FDR) was used to correct multiple comparisons when statistically significant results were obtained. SPSS (ver. 17, IBM Corp.), RStudio (ver. 4.3.0, RStudio Inc.), and GraphPad Prism (ver.10.2.2, GraphPad Software) were used to conduct these statistical tests.

For the mediation analysis^37^, which examined the mediating effects of Glx (or GABA) levels on SOL across two sleep sessions, we applied the Yeo-Johnson transformation^39^ to SOL, Glx levels, and GABA levels, using an RStudio function to improve data normality. This transformation was done because linear regressions used in the mediation analysis assume data normality. We defined the ‘Day’ factor as an independent variable (Day 1 or Day 2), the Glx (or GABA) level as a mediator, and SOL as a dependent variable. We used a tripartite model for the mediation analysis using a structural equations model to incorporate repeated measures^38^. In addition, we accounted for the repeated observations by each participant by calculating standard errors that considered the within-participant correlation. We used the Monte Carlo simulation with 500 replications to test the significance of the mediation effect because the Monte Carlo simulation does not require the assumption of normality, unlike traditional methods, including the Sobel method^37^. We used Stata Standard Edition version 18 (StataCorp.) for this analysis.

## Data availability

The datasets generated and/or analyzed in the current study will be available as source data.

## Acknowledgments

This work was supported by the NIH (R01EY031705, R01EY019466, R01EY027841), NSF (NSF-BSF: 2241417) and KAKENHI (JP20KK0268).

## Author contributions

TY and YS designed the study. TY and TLC performed the experiments. TY and AC analyzed the data. TY, TW, and YS wrote the manuscript.

