## Supplemental data for "Sleep is associated with reduction of excitatory signaling in medial prefrontal cortex"

### SUPPLEMENTARY INFORMATION

3 tables and 3 figure

**Supplementary Table 1: Sleep Structures**

| | Day 1 | | | Day 2 | | | Wilcoxon signed-rank test | |
|---|---|---|---|---|---|---|---|---|
| | | | | | | | Z | P |
| SOL (min) | 15.3 | ± | 2.99 | 9.1 | ± | 1.36 | -2.67 | 0.008 |
| WASO (min) | 13.5 | ± | 3.50 | 11.5 | ± | 3.17 | -0.56 | 0.575 |
| Stage N1 (min) | 25.4 | ± | 3.39 | 25.0 | ± | 2.60 | -0.93 | 0.926 |
| Stage N2 (min) | 27.3 | ± | 2.94 | 31.2 | ± | 3.24 | 1.66 | 0.096 |
| Stage N3 (min) | 9.5 | ± | 2.71 | 13.9 | ± | 3.67 | 1.05 | 0.293 |
| Stage R (min) | 3.3 | ± | 1.04 | 3.9 | ± | 0.91 | 0.77 | 0.441 |
| Sleep efficiency (%) | 76.1 | ± | 4.57 | 83.7 | ± | 4.12 | 1.76 | 0.079 |
| SSS score | 1.8 | ± | 0.2 | 2.0 | ± | 0.32 | 1.00 | 0.317 |

SOL, sleep-onset latency. WASO, wake after sleep onset. SSS, Stanford Sleepiness Scale. Values are means ± SEM. The related-samples Wilcoxon signed-rank test was applied to test whether each parameter was significantly different between sleep sessions (Days 1 vs. 2). SSS scores were obtained from 5 participants. See Supplementary Table 2 for the number of subjects who showed each sleep stage. When a subject missed N3 or REM sleep, we gave 0 min for that stage.

**Supplementary Table 2: Number of participants Reaching Each Sleep Stage**

| Sleep Stage | Day 1 | Day 2 | Both days |
|---|---|---|---|
| W | 20 | 20 | 20 |
| N1 | 20 | 20 | 20 |
| N2 | 20 | 20 | 20 |
| N3 | 13 | 12 | 10 |
| REM | 8 | 14 | 7 |

The column "Both days" indicates the number of participants who reached each sleep stage on both Day1 and Day2.

**Supplementary Table 3: Number of Participants in MRS Analysis for Each Sleep Stage**

| Sleep Stage | Day 1 | Day 2 | Both days |
|---|---|---|---|
| W | 20 | 20 | 20 |
| N1 | 17 | 20 | 17 |
| N2 | 20 | 20 | 20 |
| N3 | 11 | 9 | 6 |

Due to the temporal co-registration between MRS and PSG, some sleep stages were prioritized so that the numbers of data here do not necessarily match to Supplementary Table 2. The column "Both days" indicates the number of participants who reached each sleep stage with usable MRS data on both Day 1 and Day 2.

**Supplementary Figures**

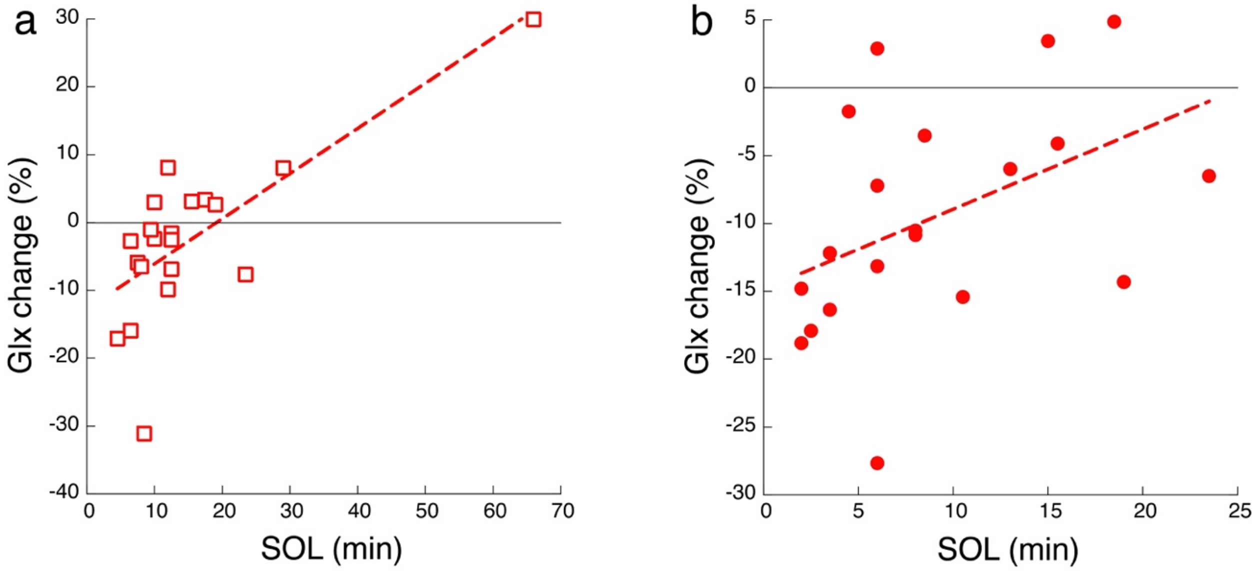


**Supplementary Figure 1. Scatter plots showing the relationship between SOL and the change in Glx levels during NREM sleep for Day 1 (a) and Day 2 (b).**

The dotted lines represent the best-fit least squares lines, included for visualization purposes only. The zero line on the Y-axis indicates the baseline Glx level, averaged during wakefulness.

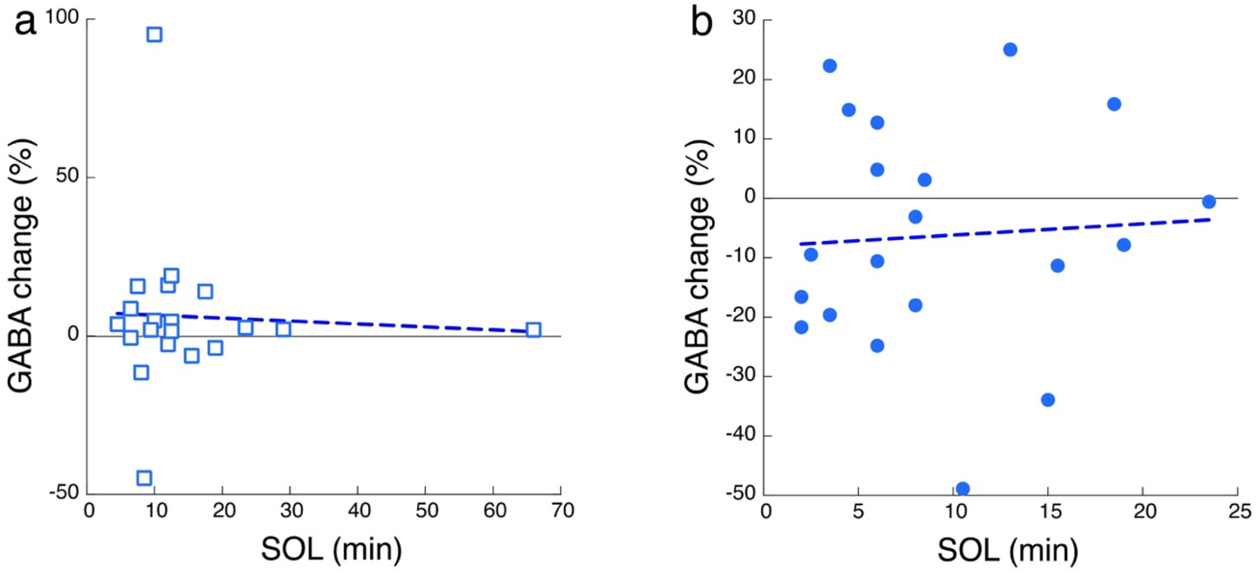


**Supplementary Figure 2. Scatter plots showing the relationship between SOL and the change in GABA levels during NREM sleep for Day 1 (a) and Day 2 (b).**

The dotted lines represent the best-fit least squares lines, included for visualization purposes only. The zero line on the Y-axis indicates the baseline GABA level, averaged during wakefulness.

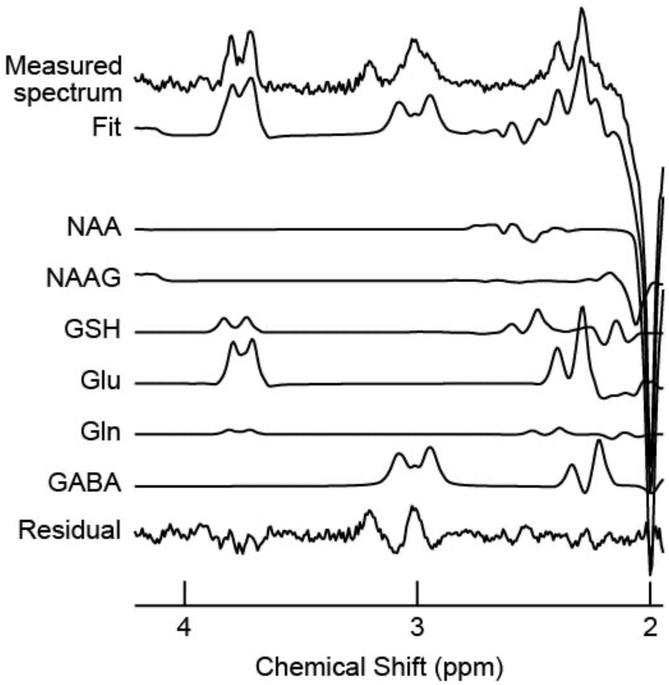


**Supplementary Figure 3. Example MRS spectra from the voxel in the mPFC.**

The top row displays the measured spectrum. The second row, labeled 'Fit', corresponds to the spectrum processed using LCModel. The bottom row, labeled 'Residual', indicates the residual spectrum after the fitting process. The remaining rows show the individual metabolite fits detectable with the specified acquisition technique. The abbreviations NAA, NAAG, GSH, Glu, Gln, and GABA represent N-acetylaspartate, N-acetylaspartylglutamate, glutathione, glutamate, glutamine, and gamma-aminobutyric acid, respectively. The term Glx refers to the sum of Glu and Gln as quantified by LCModel.
